# Oral administration of a dual ET_A_/ET_B_ receptor antagonist promotes neuroprotection in a rodent model of glaucoma

**DOI:** 10.1101/2020.10.27.357384

**Authors:** Nolan R. McGrady, Dorota L. Stankowska, Hayden B. Jefferies, Raghu R. Krishnamoorthy

## Abstract

**Purpose:** Glaucoma is a neurodegenerative disease associated with elevated intraocular pressure and characterized by optic nerve axonal degeneration, cupping of the optic disc and loss of retinal ganglion cells (RGCs). The endothelin (ET) system of vasoactive peptides (ET-1, ET-2, ET-3) and their G-protein coupled receptors (ET_A_ and ET_B_ receptors) have been shown to be contributors to the pathophysiology of glaucoma. The purpose of this study was to determine if administration of the endothelin receptor antagonist, macitentan, after the onset of IOP elevation, was neuroprotective to retinal ganglion cells in ocular hypertensive rats.

**Methods:** Brown Norway rats were subjected to the Morrison model of ocular hypertension by injection of hypertonic saline through episcleral veins. Macitentan (5 and 10 mg/kg body wt/day) was administered orally following the elevation of IOP and rats with IOP elevation were maintained for 4 weeks. RGC function was determined by pattern electroretinography at 2 and 4 weeks post IOP elevation. Rats were euthanized by approved humane methods and retinal flat-mounts generated were immunostained with RGC-selective markers RBPMS and Brn3a. RGC counts were conducted in a masked manner.

**Results:** A significant protection of retinal ganglion cells against cell loss was found following oral administration of macitentan (5 and 10 mg/kg body wt/day) in rats with elevated intraocular pressure. In addition, treatment with macitentan was able to preserve RGC function as measured by pattern ERG analysis.

**Conclusions:** Macitentan was able to promote neuroprotection independent of IOP-lowering suggesting that this could complement existing treatments to prevent neurodegeneration during ocular hypertension. The findings presented have implications for the use of macitentan as an oral formulation to promote neuroprotection in glaucoma patients.

## Introduction

Primary open-angle glaucoma (POAG) is commonly described as a complex family of optic neuropathies leading to irreversible blindness. The greatest risk factors for POAG include age and race, however, the most attributable risk factor for the development of glaucoma is an increase in intraocular pressure (IOP). To date, glaucoma therapies have solely focused on lowering IOP either surgically or pharmacologically. Although these treatments have proven effective, in some cases progression of glaucomatous degeneration still persists [1]. Given that POAG results in degeneration of the retinal ganglion cells (RGCs) which are terminally differentiated neurons that lack the ability to regenerate, it is imperative to develop approaches to promote robust neuroprotection of these cells. Due to the complex nature of the disease and the limitations of IOP-focused treatments, it would be beneficial to develop IOP-independent neuroprotective therapies. These therapies could possibly be used as an alternative to or in conjunction with current treatments. One area of interest that has emerged as a potential therapeutic target is the endothelin system.

The endothelin system is composed of three distinct 21-amino acid peptides, endothelin-1 (ET-1), endothelin-2 (ET-2) and endothelin-3 (ET-3) encoded by three separate genes in the human genome [2]. ET-1 was first described as a potent vasoactive peptide [3] and has been shown to play a significant role in vascular homeostasis [4]. Endothelin peptides bind to either of two distinct G-protein coupled receptors (GPCRs), endothelin A (ET_A_) receptor and endothelin B (ET_B_) receptor [5]. Studies revealed all three peptides have approximately equal affinity for the ET_B_ receptor however ET-3 has much lower affinity for the ET_A_ receptor making ET-3 an ET_B_ receptor selective agonist [6, 7]. Since its original discovery, the endothelin system has been found to be expressed in other areas of the body including the renal system [8–10], brain [11, 12] and eye [13].

The current study focuses on the endothelin system in the eye and within the eye ET-1 and its receptors have been shown to be expressed in multiple tissues including the iris, choroid, retina, RPE and cornea [14, 15]. Moreover, studies have shown increased levels of endonthelin-1 (ET-1) in both the aqueous humor and plasma of patients and in animal models of glaucoma [16–19]. In animal models ocular administration of ET-1 has been shown to produce ischemic damage leading to RGC axon injury and optic nerve degeneration [20–22]. Administration of 2 nmole of ET-1 produced alterations in RGC anterograde transport and disruption of the transportation of mitochondrial subcomponents [23]. Another study also showed a reversible disruption of the optic nerve fast axonal transport using low concentration ET-1 (1nM) [24]. A recent study found that mitochondrial integrity and function is important to RGC health by preserving mitochondrial activity using nicotinamide (vitamin B3) protects RGCs from glaucomatous damage [25]. Other studies have also shown the impact of mitochondrial dysfunction on RGC degeneration and as risk factors for developing POAG [26, 27].

While there might still be some debate about the initial site of damage, clinical examinations and experimental findings have provided evidence that the optic nerve head (ONH) is the first site of damage in glaucomatous degeneration [28–30]. Evidence suggests the activation of optic nerve head astrocytes (ONHAs) also contributes to axonal degeneration [31–33]. ET-1 has been shown to activate ONHAs which can be inhibited by blocking either the ET_A_ receptor or ET_B_ receptor [34]. In addition both ET_A_ and ET_B_ receptors have been shown to contribute to ET-1 mediated release of collagen I and collagen VI from lamina cribrosa cells [35]. Multiple studies have shown the significant contribution of the ET_B_ receptor to glaucomatous degeneration [36–39]. More recently, our lab showed the ET_A_ receptor’s contribution to glaucomatous degeneration following IOP-elevation [40].

ET-1 has the ability to induce an array of degenerative effects observed in POAG and studies have found in an inheritable mouse model of glaucoma that the endothelin system is activated early in the disease and occurs prior to any noticeable morphological damage [41]. These findings highlight the likelihood of the endothelin system being a potential target for neuroprotective intervention. Further evidence for this was demonstrated by the prevention of glaucomatous damage through administration of dual endothelin receptor antagonists: bosentan and macitentan [41, 42]. While these previous studies with dual endothelin receptor antagonists showed significant protection, administration of the drugs was started prior to the clinical manifestation (IOP elevation) of the disease. In contrast, our study aims to determine if treatment with macitentan started following the induction of IOP elevation can promote neuroprotection of RGCs.

## Methods

### Induction of Ocular Hypertension and IOP Measurements

Adult male retired breeder Brown Norway rats were used for all experiments in this study. All procedures involving animals were carried out in accordance with the ARVO resolution for the Use of Animals in Ophthalmic and Vision Research and approved by the University of North Texas Health Science Center (UNTHSC) Institutional Animal Care and Use Committee (IACUC). The Morrison model of ocular hypertension was carried out as previously described [40, 43]. Briefly, rats were anesthetized by intraperitoneal injection (100 μL/100 g body wt) of a ketamine (VEDCO) / xylazine (VEDCO) / acepromazine (Lloyd Laboratories) cocktail with final concentrations of 55.6 mg/mL / 5.6 mg/mL / 11.1 mg/mL, respectively. A small incision was made in the conjunctiva to expose the episcleral veins. 1.8M NaCl solution was then injected via glass micropipette (TIP01TW1F, WPI) at a flow rate of 309μL/min for 10 seconds in one eye while the fellow eye served as the contralateral control. Antibiotic ointment was applied to the surgical area to prevent any infections. Induction of ocular hypertension typically occurs 7-10 days following Morrison surgery. Macitentan treatment (5 and 10 mg/kg body wt/day) was started following the induction of IOP elevation. To ensure proper consumption, macitentan was administered orally by mixing the drug into Clear H_2_o Dietary Gel. Rats were monitored to confirm complete consumption. IOP was measured using a TonoLab tonometer (iCare, Finland) 2-3 times a week for the duration of the study. For each eye (IOP elevated and contralateral control), 10 pressure readings were recorded and averaged to give one IOP measurement. Total IOP exposure was calculated by computing the integral product of IOP elevation and the number of days for which it was maintained (expressed as mmHg-days).

### Retinal Flat Mounts and Immunostaining

Rats were euthanized by intraperitoneal injection of pentobarbital (120 mg/kg body wt). Each eye was visibly marked on the exterior of the superior part of the globe using a permanent non-washable marker to preserve orientation before eyes were enucleated. Following enucleation, eyes were briefly washed in ice-cold PBS. An ophthalmic micro surgical knife (MVR 20G, 160710, Cambrian-Medical) was used to create a small incision just posterior to the limbus and eyes were then incubated in 4% PFA for 30 min at room temperature. Small surgical scissors were then used to cut circumferentially around the globe until the anterior segment, including the lens, could be completely removed. The posterior segments were then placed in 4% PFA overnight at 4°C. Posterior segments were washed 3 times in 1 mL PBS for 10 min each wash. Posterior segments were then immersed in permeabilization buffer (0.1% sodium citrate and 0.2% Triton-X-100 in PBS) for 10 min and then washed 3 times in 1 mL PBS for 10 min each wash. Posterior segments were then immersed in blocking buffer (5% normal donkey serum and 5% bovine serum albumin) and incubated overnight at 4°C. Blocking buffer was removed and posterior segments were washed 3 times in 1 mL PBS for 10 min each wash. Posterior segments were the immersed and incubated in primary antibody solution (PBS containing 1% BSA) for 72 hours at 4°C. Following incubation with primary antibody, posterior segments were washed 3 times in 1 mL PBS for 10 min each wash. Posterior segments were then placed in secondary antibody solution (PBS containing 1% BSA) and incubated overnight at 4°C. Posterior segments were removed from the secondary antibody solution and washed 3 times in 1 mL PBS for 10 min each wash. Retinas were carefully separated from the retinal pigment epithelium and completely removed from the posterior segment. Small surgical scissors were used to make 4 cuts around the retina to allow it to be flattened. Retinas were then placed onto glass slides and mounted using Prolong^®^ Gold anti-fade reagent (P36935, Invitrogen). Primary antibodies used were goat anti-Brn3a (1:500, sc-31894, Santa Cruz) and rabbit anti-RBPMS (1:100, GTX118619, GeneTex). The corresponding secondary antibodies used were donkey anti-goat Alexa 647 (1:1000, Invitrogen) and donkey anti-rabbit Alexa 488 (1:1000, Invitrogen).

### Flat Mount Imaging and RGC Counts

All images were taken using a LSM 510 Meta confocal microscope. For each retina, two pictures were taken at each eccentricity for each of the superior, inferior, nasal and temporal quadrants. For this study, Z-stacked images were taken at eccentricities 1 and 2 which constitute 1/3 and 2/3, respectively, the width of the retina as measured from the optic nerve head. Images were randomized and RGC counts were performed by a masked observer using ImageJ software for Mac computers.

### Pattern Electroretinography

Rats were anesthetized by intraperitoneal injection (100 μL/100 g body wt) of a ketamine (VEDCO) / xylazine (VEDCO) / acepromazine (Lloyd Laboratories) cocktail with final concentrations of 55.6 mg/mL / 5.6 mg/mL / 11.1 mg/mL, respectively. Pattern ERG analysis was carried out using the Jörvec instrument (Intelligent hearing systems, Miami, FL). Rats were placed onto a heated platform adapted for rats which allowed unobstructed views of the visual stimulus monitors. Rats were maintained at 37°C for the duration of the procedure. Reference and ground electrodes were placed subcutaneously in the scalp and base of the tail, respectively. Saline eye drops were applied to both eyes to prevent drying and corneal electrodes were positioned at the lower fornix in contact with the eye globe. LED monitors were used to display contrast-reversing horizontal bars at a spatial frequency of 0.095 cycles/degree and luminance of 500cd/m^2^. Pattern ERG waveforms for each run consisted of 372 sweeps which were then averaged.

## Results

### Macitentan prevents death of RGCs following IOP elevation

To determine the effectiveness of macitentan in preventing RGC death following IOP elevation two cohorts of Brown Norway rats were subjected to IOP elevation in one eye and were maintained for 4 weeks. For both cohorts, administration of macitentan was started following the induction of IOP elevation. In the first cohort rats were given 10 mg/kg body wt/day macitentan (in gel packs) for the duration of the study. The extent of IOP exposure was then calculated for both the untreated and macitentan treated rats. No significant difference was found in the average IOP exposure between untreated (135mmHg-days) and macitentan treated (115mmHg-days) rats following macitentan treatment (Figure 1).

**Figure 1:**
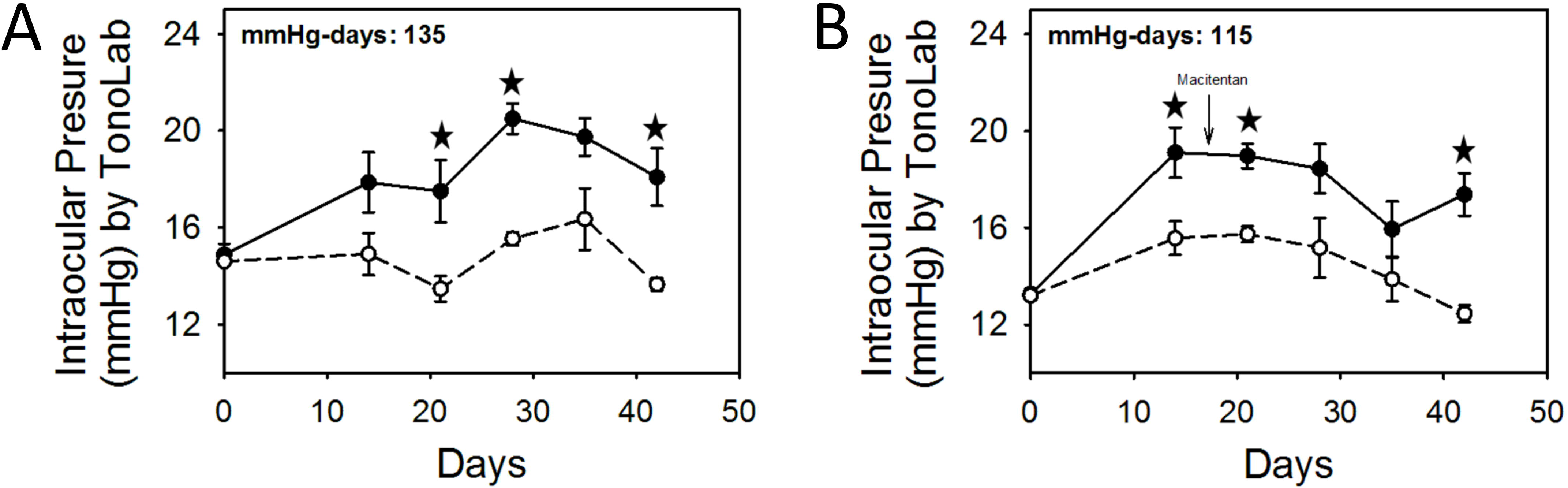
IOP profiles of Brown Norway rats with 4 weeks of IOP elevation. Plots of average IOP measurements for IOP-elevated (black circle, solid line) and contralateral eyes (white circle, dashed line) in untreated **(A)** and macitentan **(B)** fed rats. Measurements at each time point represents mean IOP ± SEM. * indicates statistical significance p<0.05 by student’s t-test, n=3.

To determine the extent of RGC loss, rats were euthanized, retinal flat mounts were prepared and immunostained with RGC selective marker RBPMS. Images were taken at eccentricities 1 and 2 (Ecc1 and Ecc2) which are 1/3^rd^ and 2/3^rd^ respectively, the width of the retina from the optic nerve head. RGC counts were performed by a masked observer and the L/R (IOP-elevated/non-elevated) ratio was determined. Following 4 weeks of IOP elevation macitentan treated rats showed significant protection against RGC loss at both eccentricities measured. At Ecc1 untreated control rats had a L/R ratio of 0.76 ± 0.15 (n=3) compared to macitentan treated rats which had a L/R ratio of 0.97 ± 0.03 (p<0.05, n=3, student’s t-test) (Figure 2). At Ecc2 untreated control rats showed a L/R ratio of 0.58 ± 0.08 (n=3) compared to macitentan treated rats which showed a L/R ratio of 0.94 ± 0.02 (p<0.05, n=3, student’s t-test) (Figure 2).

**Figure 2:**
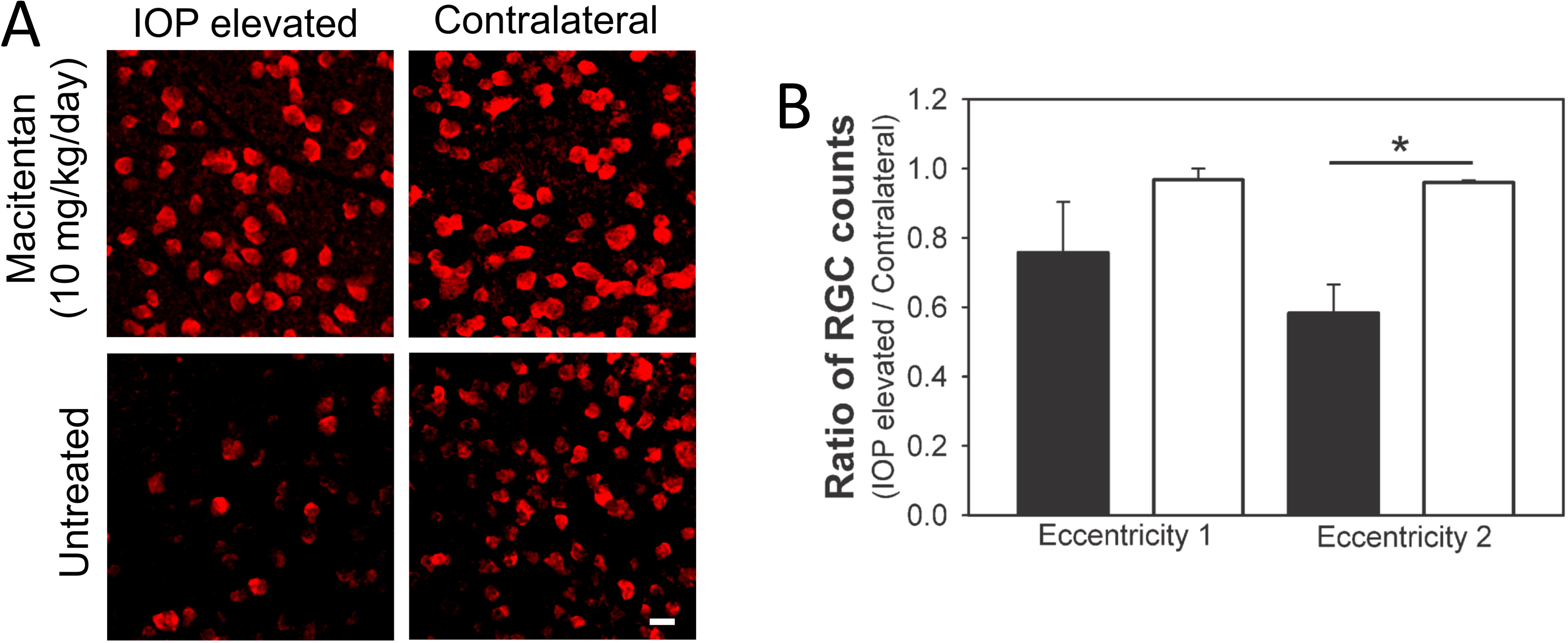
Macitentan (10 mg/kg body wt/day) enhances RGC survival in rats during IOP elevation for 4 weeks. **(A)** Representative images. RBPMS staining of RGCs in retinal flat mounts from rats which were either untreated or treated with Macitentan following IOP elevation for 1 month. Scale bar: 20 μm. **(B)** Quantification of the ratio of RBPMS-positive RGCs in IOP elevated eye (left eye) vs contralateral control eye (right eye) of Brown Norway rats for eccentricities 1 and 2, located 1/3^rd^ and 2/3^rd^ the width of the retina from the optic nerve head, respectively. Black bars represent untreated rats. White bars represent macitentan treated rats. Bars represent mean RGC count ± SEM. * indicates statistical significance, p<0.05, by student’s t-test, n=3.

In the second cohort, rats were fed 5 mg/kg body wt/day macitentan (in gel packs) three times a week. Similar to what was observed from the first cohort, there was no significant difference found in the average IOP exposures between untreated rats (171 ± 11 mmHg-days, n=3) and macitentan treated rats (172 ± 3 mmHg days, n=4) (Figure 3) indicating no effect on IOP from macitentan treatment.

**Figure 3:**
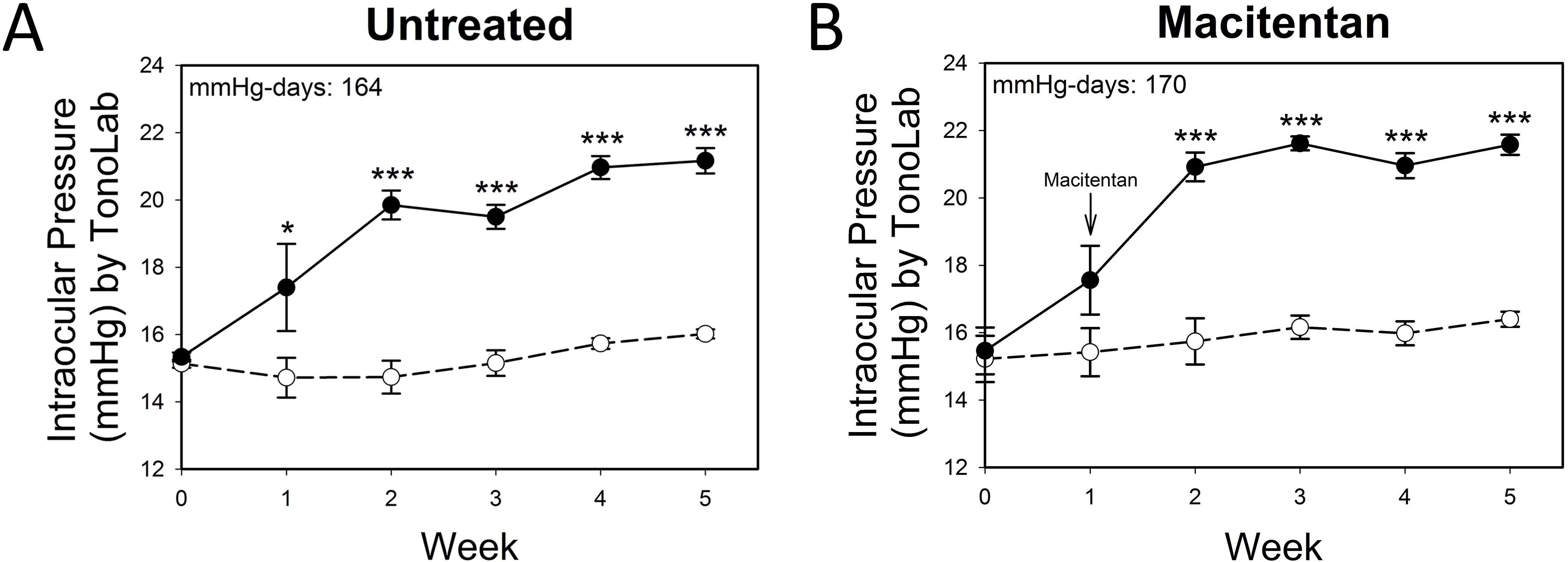
IOP profiles of Brown Norway rats with 4 weeks of IOP elevation. Plots of average IOP measurements for IOP-elevated (black circle, solid line) and contralateral eyes (white circle, dashed line) in untreated **(A)** and macitentan **(B)** fed rats. Measurements at each time point represents mean IOP ± SEM. * indicates statistical significance p<0.05 by student’s t-test, n=3.

To determine the extent of RGC loss, retinal flat mounts were prepared and immunostained with RGC selective marker Brn3a. Images were again taken at Ecc1 and Ecc2. RGC counts were performed by a masked observer and the L/R (IOP-elevated/non-elevated) ratio was determined. Following 4 weeks of IOP elevation macitentan treated rats (5 mg/kg body wt/day) showed significant protection against RGC loss in both eccentricities measured. At Ecc1 untreated control rats had a L/R ratio of 0.39 ± 0.08 (n=3) compared to macitentan treated rats which had a L/R ratio of 0.69 ± 0.11 (p<0.05, n=4, student’s t-test) (Figure 4). At Ecc2 untreated control rats showed a L/R ratio of 0.58 ± 0.06 (n=3) compared to macitentan treated rats which showed a L/R ratio of 1.06 ± 0.14 (p<0.05, n=4, student’s t-test) (Figure 4).

**Figure 4:**
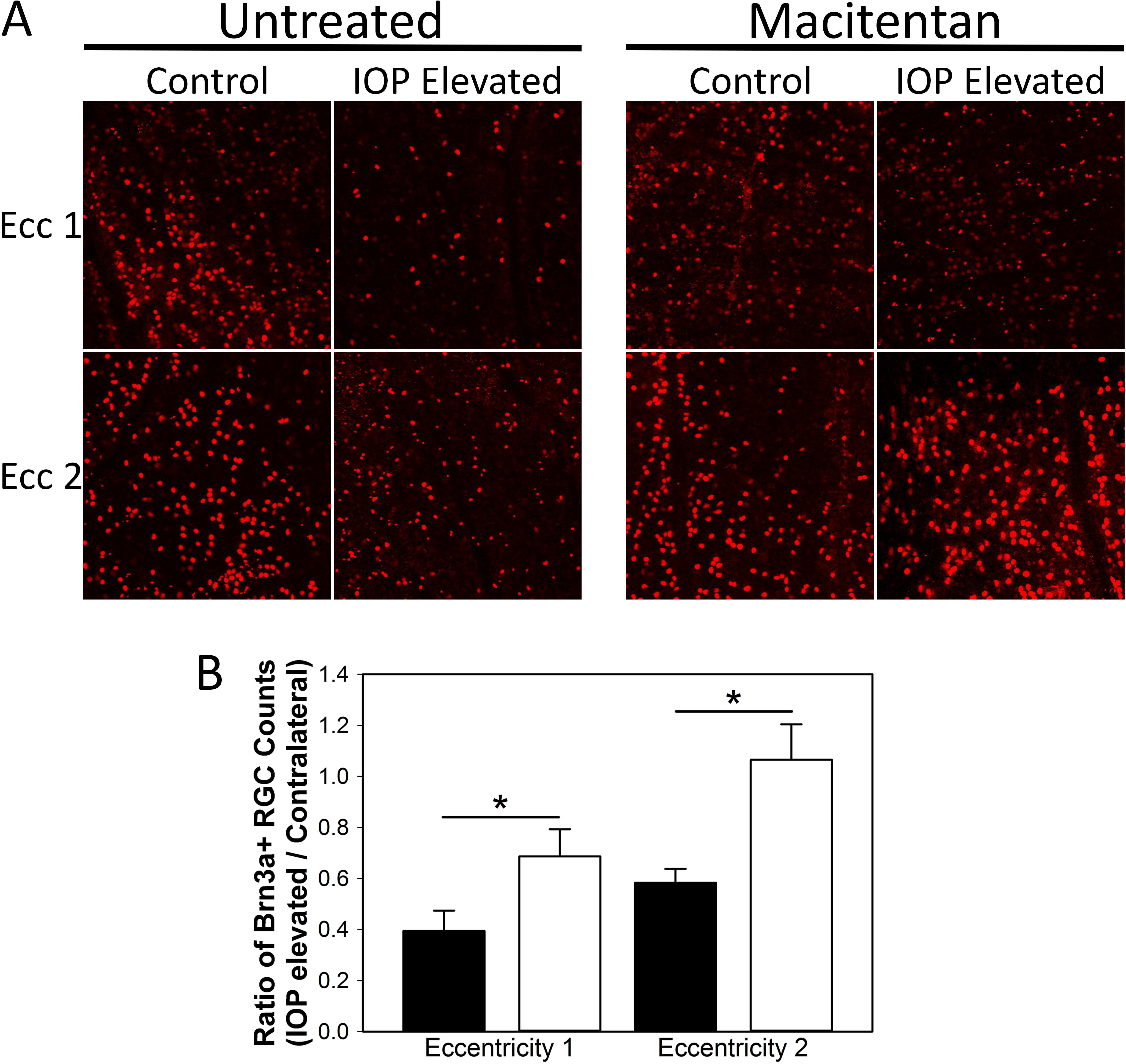
Macitentan (5 mg/kg body wt/day) enhances RGC survival in rats during IOP elevation for 4 weeks. **(A)** Representative images. Brn3a staining of RGCs in retinal flat mounts from rats which were either untreated or treated with macitentan following IOP elevation for 1 month. **(B)** Quantification of the ratio of Brn3a-positive RGCs in IOP elevated eye (left eye) vs contralateral control eye (right eye) of Brown Norway rats for eccentricities 1 and 2, located 1/3^rd^ and 2/3^rd^ the width of the retina from the optic nerve head, respectively. Black bars represent untreated rats. White bars represent macitentan treated rats. Bars represent mean RGC count ± SEM. * indicates statistical significance, p<0.05, by student’s t-test. n=3 for untreated rats, n=4 for macitentan treated rats.

### Macitentan protects RGC function following IOP elevation

In addition to RGC survival, we wanted to determine if macitentan could also protect RGC function. To visualize RGC function, pattern electroretinography (pERG) was carried out using adult retired breeder Brown Norway rats. Baseline pERGs were performed and peak latency and amplitude were recorded from naïve Brown Norway rats. The average baseline latency was measured at 87.59 ± 1.53 ms for naïve eyes (n=18) (Figure 5A). Peak amplitude at baseline was measured at 7.19 ± 0.56 μV in naïve eyes (n=18) (Figure 5B). IOP was then elevated using Morrison’s method and maintained for 2 to 4 weeks. Macitentan (5 mg/kg body wt/day) was administered after the induction of IOP. After 2 weeks of IOP elevation pERGs were recorded again. Both untreated and macitentan treated rats with 2 weeks of IOP elevation showed an increase in peak latency (untreated: 101.00 ± 6.97 ms; macitentan: 100.73 ± 5.32 ms) (Figure 5A). Pattern ERG recordings showed a reduction of peak amplitude in untreated rats while macitentan treated rats showed statistically significant preservation of peak amplitude when compared to untreated rats (untreated: 5.42 ± 0.31 μV; macitentan: 7.27 ± 0.78 μV, * p<0.05, n=5 student’s t-test) (Figure 5B). Following 4 weeks of IOP elevation pERGs were once again recorded. No difference was found in the recorded latency times between untreated and macitentan treated rats (untreated: 92.39 ± 7.19 ms; macitentan: 94.33 ± 6.35 ms) (Figure 5A). On the other hand, pERGs of macitentan fed rats (5 mg/kg body wt/day) continued to show better preservation of peak amplitude when compared to untreated rats (untreated: 5.62 ± 0.51 μV; macitentan: 6.31 ± 0.74 μV) indicating the ability of macitentan to preserve RGC function (Figure 5B).

**Figure 5:**
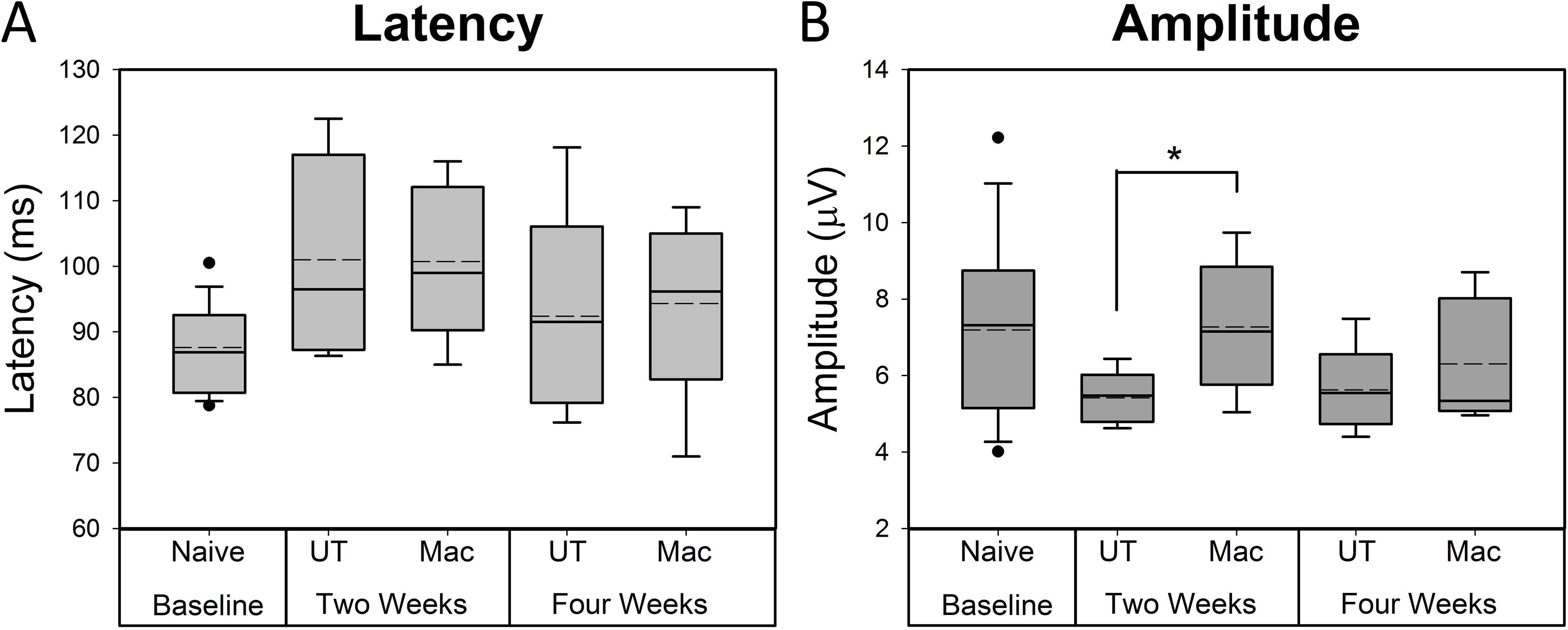
Macitentan preserves RGC function in Brown Norway rats following IOP elevation. Pattern ERG latency **(A)** and amplitude **(B)** readings of IOP elevated and control eyes for untreated and macitentan treated Brown Norway rats. Boxes represent the median, 25^th^ and 75^th^ percentiles. Whiskers represent 5^th^ and 95^th^ percentiles. Black circles represent data points outside the 5 and 95 percentiles. Dashed lines represent mean values. * indicates statistical significance, p<0.05, n=5, student’s t-test. Abbreviations: UT-untreated, Mac-macitentan treated.

## Discussion

Primary open-angle glaucoma (POAG) is a multifaceted disease however therapies for POAG only target one risk factor of the disease, elevated IOP, which is not observed in every patient (normotension). Although current treatments have proven effective for both ocular hypertensive and normotensive patients, there is a limitation on how far a patient’s IOP can be lowered. In the clinical setting, in most patients IOP cannot be lowered beyond 20% using a combination of therapies targeting aqueous humor formation and outflow [44]. Thus, it would be advantageous to develop IOP-independent alternative strategies to prevent neurodegeneration of RGCs and promote neuroprotection.

The endothelin system has been established as an important contributor to the pathophysiology of POAG [13, 45–49]. It is currently understood that there are several key cellular events that occur in the optic nerve head as well as the retina during the progression of POAG prior to RGC death [30, 50, 51]. The events include the disruption of RGC axonal transport (52-57), activation and redistribution of optic nerve head astrocytes [58–61] and changes in the optic nerve head extracellular matrix milieu [62–64]. Multiple studies have shown the ability of the endothelin system, namely ET-1, to induce these key glaucomatous events including a decrease in axonal transport [23, 24, 65, 66], proliferation of optic nerve head astrocytes [34, 67, 68] and extracellular deposition of collagens by lamina cribrosa (LC) cells [35]. Since endothelins act through either the ET_A_ and ET_B_ receptor or through both these receptors to mediate these deleterious effects, it would be prudent to use an ET_A_/ET_B_ dual receptor antagonist to block neurodegenerative effects in glaucoma. In fact, Howell et al. (2011) demonstrated that a dual-endothelin receptor antagonist, bosentan, could prevent glaucomatous neurodegeneration in the DBA/2J mouse [41]. In a subsequent study, Howell et al., (2014) used macitentan (30 mg/kg) (which has higher affinity and longer receptor occupancy times) and found it to significantly prevent axonal loss in the DBA/2J mice [42].

Macitentan has been approved by the FDA in 2013 for the treatment of pulmonary arterial hypertension and its safety in humans as already been established. In the current study we demonstrate the ability of macitentan to promote protection of RGCs following IOP elevation without lowering IOP in the Morrison’s rat model of glaucoma. Moreover, in this study, we started the macitentan treatment after the onset of IOP elevation, which simulates the clinical scenario where patients start the treatment after diagnosis of glaucoma. Considering all current clinical therapies only target IOP, an IOP-independent therapy could be a viable option for primary open-angle glaucoma patients, especially for those who continue to show glaucomatous progression. In addition to promoting RGC survival we have also demonstrated the ability of a dual endothelin receptor antagonist, macitentan, to preserve RGC function as measured by pattern electroretinography. Chou et al. (2013) demonstrated that retrograde signaling is required for pattern ERG response [69] suggesting that signal transduction between the retina and the brain was preserved in macitentan fed rats in the current study.

While we have assessed the effect of macitentan in the posterior segment of the eye, it is plausible that it could also have some beneficial effects on aqueous humor dynamics in the anterior segment of the eye (which is not obvious in the Morrison’s model due to irreversible sclerosis of the trabecular meshwork). For instance, it is understood that IOP elevation results from cellular alterations within the trabecular meshwork (TM). The alterations in the TM reduce the outflow of aqueous humor from the anterior segment of the eye which results in an increase in IOP [70]. It has been shown that the aqueous humor of patients with glaucoma show increased levels of TGF-β [71–76] which has been shown to contribute to the pathogenesis of POAG [77]. Levels of ET-1 have been shown to be increased following overexpression of TGF-β in the TM [78]. Interestingly, ET-1 was found to contribute to TGF-β induced fibrosis in skin and lung tissues which was ameliorated by bosentan [79, 80]. Macitentan was unable to prevent IOP elevation in the DBA/2J mouse model although IOP increase in this model is due to dispersion of iris pigmentation which impedes aqueous humor outflow from the trabecular meshwork. While studies that focus on this relationship in the eye are limited, it would be worth investigating the potential of a dual endothelin receptor antagonist like macitentan to prevent increased ECM deposition within the TM and possibly the LC region of the optic nerve head.

In the current study, oral administration of the dual ET_A_/ET_B_ receptor antagonist, macitentan, was found have neuroprotective effects on RGC survival at two different doses (5 and 10 mg/kg body wt/day) in ocular hypertensive rats. In addition, macitentan was able to preserve RGC function as determined by pERG analysis. The study brings up exciting possibilities for use of macitentan as an oral formulation to promote neuroprotection in glaucoma patients. This is particularly helpful to elderly patients due to non-compliance issues stemming from difficulties in installing eye drops, for instance, in patients that have motor deficits. Future studies will examine cellular mechanisms and signaling pathways contributing to macitentan-mediated neuroprotection in glaucoma.

## List of Abbreviations

POAG: Primary Open Angle Glaucoma
IOP: Intraocular Pressure
ET_A_: Endothelin Receptor A
ET_B_: Endothelin Receptor B
ET-1: Endothelin-1
pERG: Pattern Electroretinography

## Acknowledgements

The authors thank Dr. Thomas Yorio for several useful discussions and feedback on this project. The authors thank Dr. Vittorio Porciatti for training to do pattern ERG in his laboratory.

RRK and DLS performed Morrison surgeries. NRM, DLS, and HBJ measured IOP. NRM and DLS performed immunohistochemistry, imaging and pattern ERG. RRK and HBJ performed RGC counts. NRM wrote the manuscript. RRK conceived the experimental plan and designed the study.

This work was supported by an extramural grant to RRK from the National Eye Institute [EY019952] and from an intramural grant from the UNT Health Science Center. The first author, Mr. Nolan McGrady, was supported by a NIH training grant [T32 AG 020494] awarded to the Neurobiology of Ageing training program.

